# Expression of *mecA* reduces the daptomycin susceptibility of *Staphylococcus aureus*

**DOI:** 10.1101/2025.02.10.637576

**Authors:** Elizabeth V. K. Ledger, Mario Recker, Ruth C. Massey

## Abstract

*Staphylococcus aureus* is a leading cause of bacteraemia and infections caused by methicillin-resistant strains (MRSA) are especially challenging to treat. MRSA strains are resistant to front-line beta-lactams due to PBP2a, a low-affinity penicillin-binding protein encoded by *mecA*. Daptomycin is used to treat MRSA infections but is not always effective and is associated with high rates of morbidity and mortality. Understanding why daptomycin fails is crucial to devising strategies to improve treatment outcomes. Here, using a panel of clinical bacteraemia isolates, we show that MRSA strains are less susceptible to daptomycin than methicillin-susceptible (MSSA) strains. This difference in susceptibility was due to *mecA* and was independent of any changes in surface properties previously associated with altered daptomycin susceptibility. Instead, using a *mecA* transposon mutant, we found that a lack of this gene led to higher activity of the Agr quorum sensing system, resulting in an increased release of the phenol-soluble modulin toxins. Increased levels of these surfactant-like toxins prevented daptomycin from being inactivated by lipids released by the bacteria, leading to enhanced susceptibility to the antibiotic. Additionally, the clinical MRSA strains tested produced lower levels of toxins than the MSSA strains and inactivated daptomycin to a greater extent, explaining their reduced susceptibility. Expression of *mecA* in clinical MSSA strains reduced toxin production, increasing daptomycin inactivation and thereby enhancing survival. Together, these results demonstrate that *mecA* does not only affect beta-lactam susceptibility but also compromises the efficacy of the last resort antibiotic daptomycin.

## Introduction

*Staphylococcus aureus* is a major human pathogen, causing a range of infections from minor skin and soft tissue infections to life-threatening diseases like bacteraemia and endocarditis^1^. It is the most common causative agent of skin infections and is the leading Gram-positive cause of bacteraemia, responsible for over 14,000 cases in 2023 in the UK^2^. Bacteraemia requires urgent intravenous antibiotics to sterilise the bloodstream as untreated it can lead to the development of secondary infections, including deep tissue abscesses and endocarditis^3^.

The treatment prescribed depends on the antibiotic susceptibility of the strain causing the infection, with methicillin-susceptible *S. aureus* (MSSA) treated with front-line beta-lactams such as flucloxacillin or cefazolin^4,5^. Treatment options for methicillin-resistant *S. aureus* (MRSA) are more limited but include vancomycin and daptomycin^4,5^. Regardless of the susceptibility of the pathogen, long-term IV antibiotics are needed, with a minimum of a 6-week course recommended in cases of complicated bacteraemia^6^. However, despite treatment, *S. aureus* bacteraemia has a fatality rate of approximately 25%^7^.

Vancomycin can be challenging to dose correctly as it can require labour-intensive monitoring to ensure the correct serum concentrations are achieved^8^. These limitations mean that daptomycin is preferred in some cases. Daptomycin is a lipopeptide antibiotic which binds to membrane phosphatidylglycerol in a calcium-dependent manner^9,10^. This disrupts membrane integrity and cell wall synthesis, leading to cell death^11–13^. Daptomycin is rapidly bactericidal *in vitro* and resistance is extremely rare, with over 99.9% of infections caused by daptomycin-susceptible strains^14^. However, despite this, daptomycin suffers from high rates of treatment failure^15,16^. Understanding the reasons behind this is crucial to improving patient outcomes.

A range of approaches have been used to identify factors that affect daptomycin susceptibility, including studying paired clinical isolates from before and after daptomycin therapy^17–19^ and generating resistant mutants *in vitro*^*20*^. Together, these have found that daptomycin susceptibility can be influenced by changes in the cell membrane, such as an increased positive charge resulting from increased lysylphosphatidylglycerol content or altered fatty acid compositions resulting in changes to membrane fluidity^21–24^. Increases in cell wall thickness and wall teichoic acid (WTA) content have also been found to reduce daptomycin susceptibility^17,25^. Daptomycin tolerance is also thought to contribute to treatment failure. This tolerance can be phenotypic and induced by the host environment^26,27^, or due to mutations in genes such as those encoding the PitA phosphate transporter^28^ or the alkaline shock protein Asp23^29^. Finally, *S. aureus* can survive exposure to daptomycin through releasing membrane phospholipids^30^. The released phosphatidylglycerol binds to daptomycin, sequestering it and preventing it from killing the bacteria^30^. However, this interaction between lipids and daptomycin is compromised by the release of phenol-soluble modulins (PSMs), small surfactant-like toxins whose release is controlled by the accessory gene regulator (Agr) quorum sensing system^30^.

However, while much is known about some factors affecting daptomycin susceptibility, the exact mechanisms used by *S. aureus* to survive daptomycin exposure remain to be fully understood. Therefore, here we aimed to discover novel genes that affect daptomycin susceptibility with the aim of identifying therapeutic targets to improve daptomycin treatment efficacy.

## Results

### MRSA strains are less susceptible to daptomycin than MSSA strains

As a first step to identifying factors that affect the susceptibility of *S. aureus* to daptomycin, we studied a collection of 300 clinical *S. aureus* bacteraemia isolates belonging to either clonal complex (CC) 22 or 30 (CC22, *n* = 135; CC30, *n* = 165) which contained both MRSA and MSSA isolates^31^. These strains were isolated from a single hospital in the UK between 2006 and 2012. An overnight culture of each strain was adjusted to 1 × 10^8^ CFU ml^-1^ and exposed to 10 μg ml^-1^ daptomycin (corresponding to mean serum levels achieved in patients^32^) for 2 h and survival measured by colony forming unit (CFU) counts.

Daptomycin resistance is rare^14^ and so, as expected, none of the strains were able to grow in the presence of the antibiotic (Fig. 1A). However, the susceptibility of the strains varied, with a > 3-log difference in survival between the most and least susceptible isolates. There were differences in susceptibility between the clonal complexes, with the median susceptibility of the isolates from CC22 and CC30 being 4 × 10^5^ and 1 × 10^6^ CFU ml^-1^, respectively (Fig. 1A). Additionally, there was a significant difference in susceptibility between MRSA and MSSA strains, where the MRSA isolates were less susceptible to daptomycin relative to the MSSA isolates (2.2 and 5.8-fold in CC22 and CC30, respectively) (Fig. 1B – C).

**Fig. 1.**
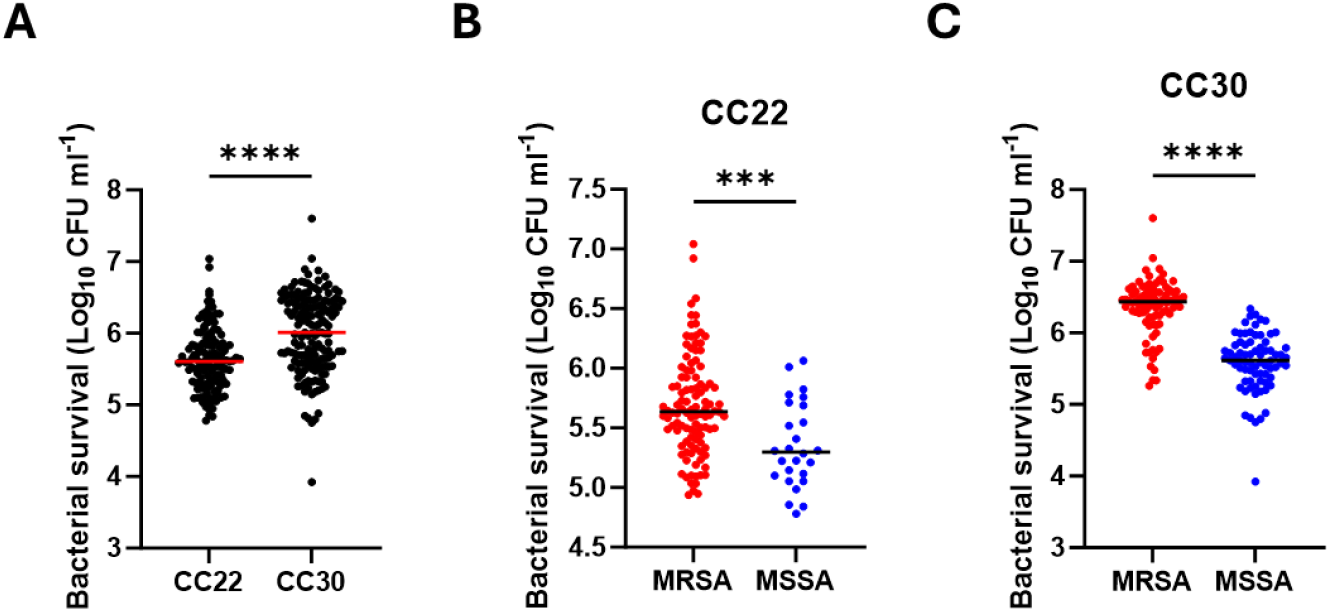
MRSA strains are less susceptible to daptomycin than MSSA strains. (**A**) Clinical isolates of *S. aureus* from CC22 (n=135) or from CC30 (n=165) at 10^8^ CFU ml^-1^ were exposed to 10 μg ml^-1^ daptomycin for 2 h before survival was determined by CFU counts. (**B-C**) Data from panel **A** are re-plotted with strains from (**B**) CC22 and (**C**) CC30 separated based on susceptibility to methicillin. Each data point represents the mean survival of three independent repeats of one strain and the median of all the strains is indicated. Data were analysed by an unpaired t-test (***, P ≤ 0.001. ****, P ≤ 0.0001).

### Expression of *mecA* reduces susceptibility of JE2 and clinical MSSA isolates to daptomycin

Daptomycin and beta-lactam resistance are typically negatively correlated, with resistance to daptomycin sensitising strains to beta-lactams, a phenomenon known as the ‘seesaw effect’^33,34^. Therefore, we aimed to understand why this negative correlation was not observed here, and why MRSA strains were less susceptible to daptomycin than MSSA strains. Firstly, we investigated whether the difference in daptomycin susceptibility between MRSA and MSSA strains was due to the presence of the *mecA* gene in MRSA strains, which encodes the low-affinity penicillin-binding protein (PBP), PBP2a. To do this, we exposed *S. aureus* JE2 wildtype (WT) and the *mecA*::Tn mutant from the Nebraska Transposon mutant library (NTML)^35^ to daptomycin for 6 h and measured survival over time by CFU counts. This demonstrated that the *mecA*::Tn mutant was approximately 100-fold more susceptible to the antibiotic than the WT strain (Fig. 2A). To confirm that this was due to *mecA* and not polar effects of the transposon insertion, we complemented the *mecA*::Tn mutant with a WT copy of *mecA* under the control of the tetracycline-inducible promoter on p*itet*, a plasmid which integrates into the genome at the *geh* locus^36^. Complementation of the *mecA*::Tn mutant with this plasmid (p*mecA*) restored daptomycin susceptibility to WT levels while complementation with the empty vector control did not (Fig. 2A). Therefore, loss of *mecA* sensitises *S. aureus* to daptomycin.

**Fig. 2.**
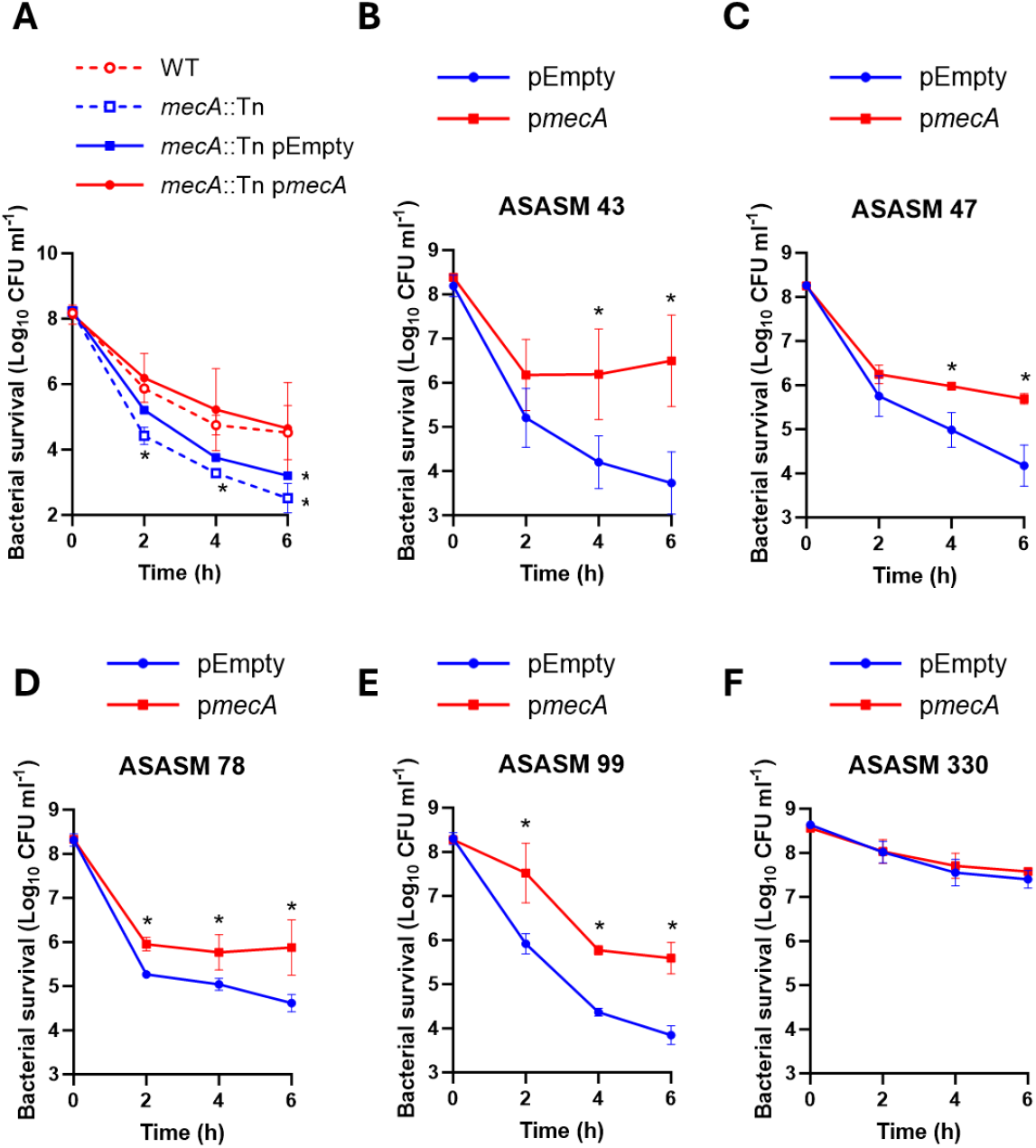
Expression of *mecA* reduces susceptibility of JE2 and clinical MSSA isolates to daptomycin. **(A)** Log_10_ CFU ml^-1^ of JE2 WT, *mecA*::Tn, *mecA*::Tn complemented with an empty plasmid or *mecA*::Tn complemented with p*mecA* during a 6 h exposure to 10 μg ml^-1^ daptomycin. Log_10_ CFU ml^-1^ of (**B**) ASASM 43, (**C**) ASASM 47, (**D**) ASASM 78, (**E**) ASASM 99 and (**F**) ASASM 330 complemented with either the empty plasmid (blue) or p*mecA* (red) during a 6 h exposure to 10 μg ml^-1^ daptomycin. Data represent the mean ± standard deviation of three independent biological repeats. Data were analysed by two-way ANOVA with Sidak’s *post-hoc* test. *, P < 0.05 (WT vs mutants (A) or pEmpty vs p*mecA* (B-F) at indicated time-points).

Next, we aimed to investigate whether expression of *mecA* also affected daptomycin susceptibility in the clinical isolates. To do this, we transformed the empty plasmid or p*mecA* into five representative MSSA strains from CC30 which spanned the observed range of daptomycin susceptibilities. As a larger difference in daptomycin susceptibility was observed between MRSA and MSSA strains from CC30 than CC22, we chose to characterise the mechanism behind this phenotype using strains from this clonal complex.

Daptomycin was rapidly bactericidal against four out of five of these strains containing the empty plasmid, causing between 3 and 4 logs of killing by 6 h (Fig. 2B – E). In each of these strains, induction of *mecA* expression led to a significant reduction in antibiotic killing (Fig. 2B – E). The fifth strain tested, ASASM330, was less susceptible than the others, with only 1 log of killing observed by 6 h and, in this case, induction of *mecA* expression had no effect on daptomycin susceptibility (Fig. 2F).

Taken together, *mecA* affects daptomycin susceptibility in both the laboratory strain JE2 and clinical isolates, with a lack of *mecA* expression increasing susceptibility and increased *mecA* expression resulting in reduced susceptibility.

### PBP2a does not affect daptomycin susceptibility by altering surface properties

The next objective was to understand why expression of *mecA* affected daptomycin susceptibility. Many factors have been identified to impact daptomycin susceptibility, including increased cell wall thickness, alterations to the cell membrane and increased positive charge on the cell surface^10^. Therefore, we next tested whether any of these properties differed between five representative MRSA and MSSA strains and whether these differences depended on *mecA*.

Firstly, we examined surface charge, using a fluorescently labelled positively charged molecule, poly-L-lysine (FITC-PLL). This demonstrated that the MRSA strains were more positively charged than the MSSA strains tested and so would be expected to show increased repulsion of the positively-charged antibiotic (Fig. 3A). However, the difference in surface charge was not dependent on PBP2a, with a *mecA*::Tn mutant having the same surface charge as the WT strain (Fig. 3A). Similarly, we observed differences in membrane fluidity, staphyloxanthin content and WTA levels between the MRSA and MSSA strains but none of these differences were dependent on *mecA* (Fig. 3B – D). There were no differences observed in cell wall thickness between the MSSA and MRSA strains or between WT and the *mecA*::Tn mutant (Fig. 3E).

**Fig. 3.**
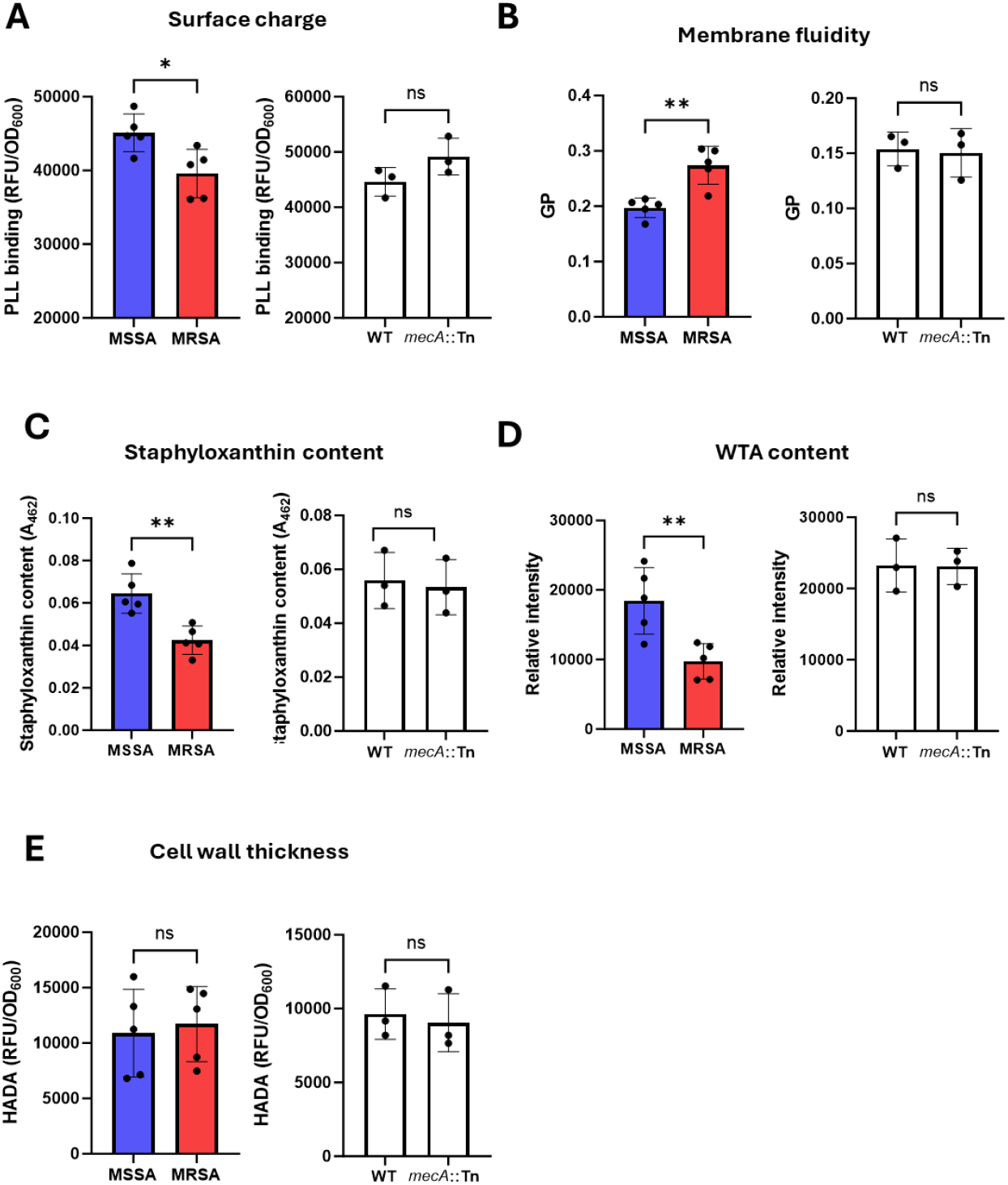
PBP2a does not affect daptomycin susceptibility by altering surface properties. **(A)** Surface charge of five MSSA and five MRSA strains (left) and JE2 WT and the *mecA*::Tn mutant (right) as determined by binding of FITC-labelled PLL. Fluorescence values were divided by OD_600_ to normalise for difference in cell density. (**B**) Membrane fluidity of five MSSA and five MRSA strains (left) and JE2 WT and the *mecA*::Tn mutant (right) as determined by the generalised polarisation index obtained by measuring fluorescence of laurdan. (**C**) Staphyloxanthin content of five MSSA and five MRSA strains (left) and JE2 WT and the *mecA*::Tn mutant (right). (**D**) Wall teichoic acid content of five MSSA and five MRSA strains (left) and JE2 WT and the *mecA*::Tn mutant (right) as determined by WTA extraction, analysis by native PAGE and visualisation with alcian blue. Intensity was quantified using ImageJ. (**E**) Cell wall thickness of five MSSA and five MRSA strains (left) and JE2 WT and the *mecA*::Tn mutant (right) as determined by the incorporation of the fluorescent D-amino acid analogue HADA. Fluorescence values were divided by OD_600_ to normalise for difference in cell density. In the left hand graph of each panel, each data point represents the mean of three independent repeats of one strain. In the right hand graph of each panel, data represent the mean ± standard deviation of three independent repeats. Data were analysed by t-test. *, P ≤ 0.05. **, P ≤ 0.01.

Therefore, the *mecA*-dependent reduced daptomycin susceptibility of MRSA strains compared to MSSA could not be explained by differences in cell surface charge, membrane fluidity, staphyloxanthin levels, WTA content or cell wall thickness.

### Expression of *mecA* reduces daptomycin susceptibility by increasing inactivation of the antibiotic

Susceptibility to daptomycin has previously been linked to activity of the Agr quorum sensing system, with Δ*agr* mutants showing reduced susceptibility compared to the WT strain^30^. This is because *S. aureus* releases phospholipids which sequester daptomycin, inactivating it. This binding of phospholipids to daptomycin is compromised by the production of PSMs, small surfactant-like toxins whose expression is controlled by Agr^30^. Therefore, next we determined whether this could explain the observed difference in daptomycin susceptibility between MRSA and MSSA strains.

Firstly, we measured whether there were differences in the degree to which daptomycin was inactivated by the MRSA strains versus the MSSA strains. To do this, the strains were exposed to daptomycin for 6 h before the activity of daptomycin remaining in the supernatant was quantified using a zone of inhibition assay. This revealed that all five of the MRSA strains fully inactivated daptomycin, whereas the MSSA strains inactivated daptomycin to a lesser degree, with an average of 40 % of the starting activity of daptomycin remaining in the supernatant after 6 h (Fig. 4A). The ability of *S. aureus* to inactivate daptomycin was dependent on *mecA*, with a *mecA*::Tn mutant inactivating significantly less daptomycin over 6 h than the WT strain (Fig. 4B). Complementation of the *mecA*::Tn mutant with a plasmid expressing *mecA*, but not with an empty plasmid control, restored daptomycin inactivation to WT levels (Fig. 4B).

**Fig. 4.**
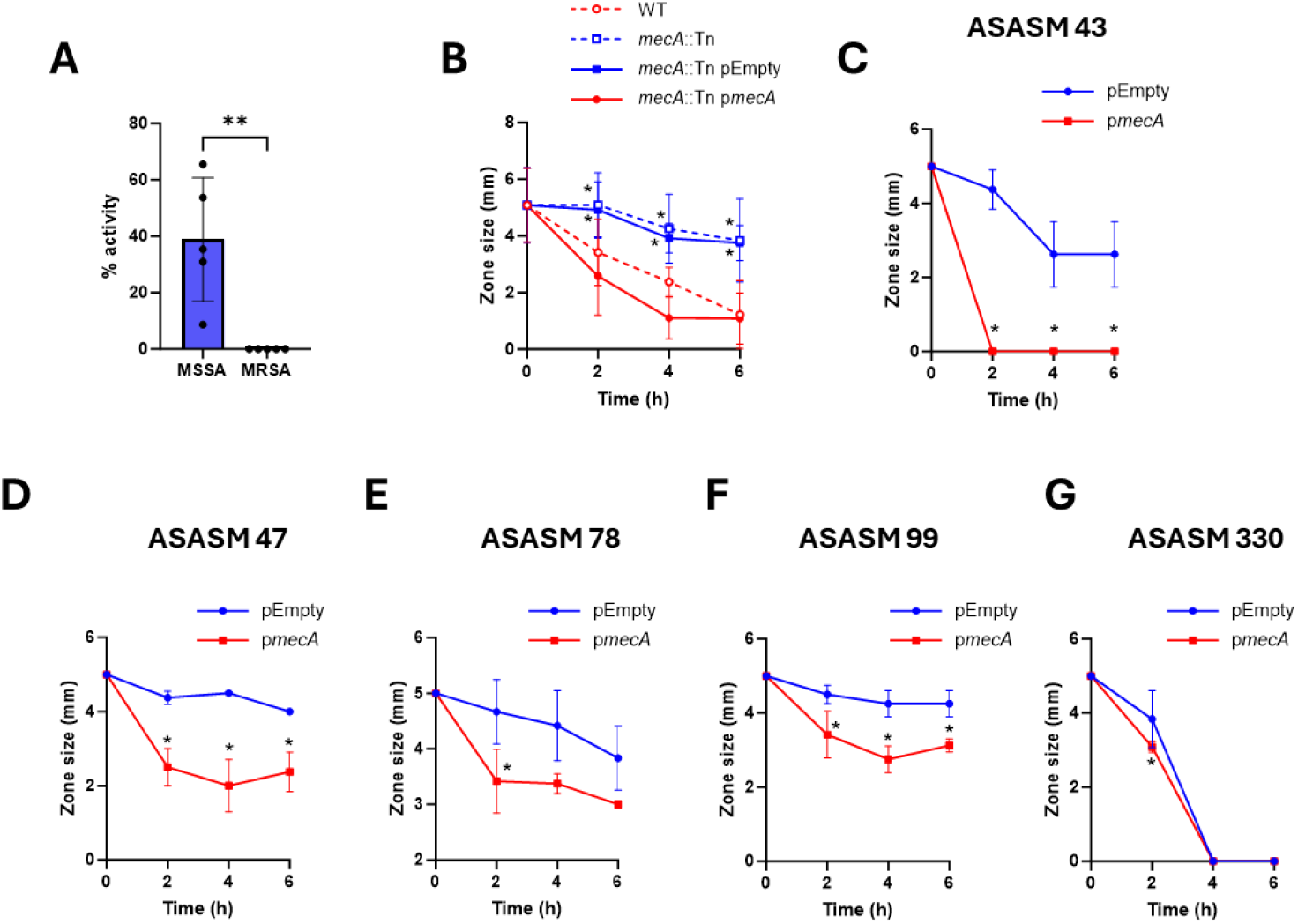
Expression of *mecA* reduces daptomycin susceptibility by increasing inactivation of the antibiotic. (**A**) Percentage of daptomycin activity remaining in the supernatant as measured by a zone of inhibition assay after exposure of five MSSA strains and five MRSA strains to 10 μg ml^-1^daptomycin for 6 h. (**B**) Daptomycin activity remaining in the supernatant after exposure of JE2 WT, *mecA*::Tn, *mecA*::Tn complemented with an empty plasmid or *mecA*::Tn complemented with p*mecA* to 10 μg ml^-1^ daptomycin. Daptomycin activity remaining in the supernatant after exposure of (**C**) ASASM 43, (**D**) ASASM 47, (**E**) ASASM 78, (**F**) ASASM 99 and (**G**) ASASM 330 complemented with either the empty plasmid (blue) or p*mecA* (red) to 10 μg ml^-1^ daptomycin. In panel A, each data point represents the mean of three independent repeats of one strain and data were analysed by t-test. All other graphs represent the mean ± standard deviation of three independent repeats. Data were analysed by two-way ANOVA with Sidak’s *post-hoc* test. *, P ≤ 0.05, **, P ≤ 0.01 (MRSA vs MSSA in panel A; WT vs other strains at indicated time-points in panel B; pEmpty vs p*mecA* at indicated time-points in panels C-G).

Next, we investigated whether induction of *mecA* expression in the MSSA strains affected their ability to inactivate daptomycin. In line with the daptomycin susceptibility data in Fig. 2B – F, expression of *mecA* in each of the MSSA strains except ASASM330 led to increased daptomycin inactivation compared to the empty plasmid control (Fig. 4C – G). ASASM330 fully inactivated daptomycin in the absence of *mecA*, explaining the low susceptibility to daptomycin observed in Fig. 2F (Fig. 4G)

Taken together, the expression of *mecA* increases the ability of *S. aureus* to inactivate daptomycin, resulting in reduced susceptibility to the antibiotic.

### Expression of *mecA* increases daptomycin inactivation by reducing production of PSMs

The final objective was to determine why expression of *mecA* affected daptomycin inactivation. Previous work has demonstrated that methicillin resistance lowers Agr activity^37,38^, and as such, we tested whether the effect on daptomycin susceptibility reported here was also due to the expression of *mecA* suppressing Agr activity and the resulting production of PSMs. Firstly, we measured the haemolytic ability of the strains as a proxy for Agr activity. This demonstrated that, as expected, the MSSA strains were more haemolytic than the MRSA strains, indicating that they had higher levels of Agr activity (Fig. 5A). Additionally, this was dependent on *mecA*, as the *mecA*::Tn mutant was more haemolytic than the WT strain (Fig. 5B) and complementation of the MSSA strains with p*mecA* led to a reduction in haemolysis compared to the empty plasmid controls (Fig. 5C).

**Fig. 5.**
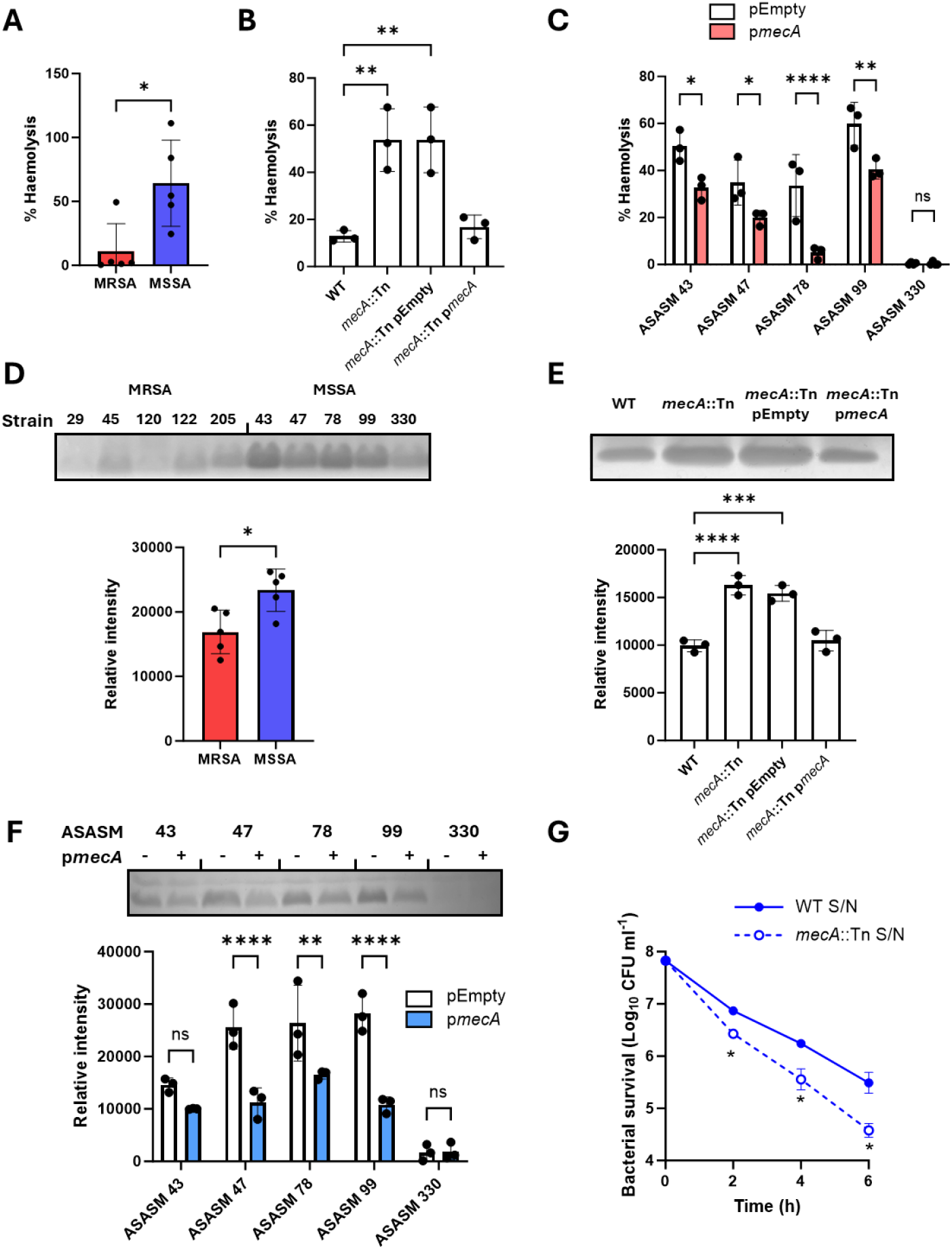
Expression of *mecA* increases daptomycin inactivation by reducing production of PSMs. (**A**) Percent haemolysis of sheep erythrocytes by supernatants from five MRSA and five MSSA strains. (**B**) Percent haemolysis caused by supernatants from JE2 WT, *mecA*::Tn and *mecA*::Tn complemented with the empty plasmid or with p*mecA*. (**C**) Percent haemolysis of supernatants from five MSSA strains complemented with either the empty plasmid or with p*mecA*. (**D**) Supernatants from five MRSA and five MSSA strains were analysed by SDS-PAGE to visualise and quantify the PSMs. (**E**) Supernatants from JE2 WT, *mecA*::Tn and *mecA*::Tn complemented with the empty plasmid or with p*mecA* were analysed by SDS-PAGE to visualise and quantify the PSMs. (**F**) Supernatants from five MSSA strains complemented with either the empty plasmid or with p*mecA* were analysed by DS-PAGE to visualise and quantify the PSMs. (**G**) Log_10_ CFU ml^-1^ of JE2 WT during a 6 h exposure to 10 μg ml^-1^ daptomycin in spent culture supernatant from either the WT strain or from the *mecA*::Tn mutant. In panels A and D, each data point represents the mean of three independent repeats of one strain and data were analysed by t-test. Data in B, C, E, F and G represent the mean ± standard deviation of three independent repeats. Data in B and E were analysed by one-way ANOVA with Dunnett’s *post-hoc* test. Data in C, F and G were analysed by two-way ANOVA with Sidak’s *post-hoc* test. *, P ≤ 0.05; ** P ≤ 0.01; *** P ≤ 0.001; ****P ≤ 0.0001. In panels D, E and F, representative images are shown from three repeats.

Following on from this, we visualised the amount of PSMs released into the supernatant using SDS-PAGE, where PSMs (molecular weight 2 – 5 kDa) migrate as a band ahead of the bromophenol blue dye front^39^. As expected, this agreed with the haemolysis data, with the MSSA strains producing more PSMs than MRSA strains (Fig. 5D), the *mecA*::Tn mutant producing more PSMs than the WT strain (Fig. 5E) and complementation of the MSSA strains with p*mecA* leading to a decrease in PSM production compared to the empty plasmid controls (Fig. 5F).

As a final confirmation that the reduced daptomycin inactivation and subsequent increased daptomycin susceptibility were due to the release of PSMs, the JE2 WT strain (which produces relatively lower amounts of PSMs) was exposed to daptomycin in spent culture supernatant from either the WT strain or the *mecA*::Tn mutant strain (which produces relatively higher levels of PSMs). The WT strain was more susceptible to daptomycin when antibiotic exposure occurred in the presence of supernatant from the *mecA*::Tn mutant compared to supernatant from the WT strain, with over a 9-fold difference in susceptibility observed at 6 h (Fig. 5G).

Taken together, the expression of *mecA* reduces Agr activity and the associated PSM production, enabling daptomycin to be inactivated by the released phospholipids and promoting bacterial survival.

## Discussion

*S. aureus* is a leading cause of bacteraemia and when the infection is caused by an MRSA strain, daptomycin is one of the few recommended treatment options. Unfortunately, daptomycin treatment is not always effective, failing in 20 – 30% of cases and resulting in high mortality rates^16,40^. Understanding why this treatment failure occurs is crucial to developing improved approaches to manage staphylococcal infections. Here, by testing the daptomycin susceptibility of 300 clinical bacteraemia isolates, we identified *mecA* as a crucial determinant of daptomycin susceptibility.

The *mecA* gene encodes the low affinity PBP, PBP2a and confers resistance to beta-lactam antibiotics. It is encoded within the SCC*mec* element, a region of variable size which can be excised and integrated into genomes using the CcrAB recombinases^41^. The size of the SCC*mec* element varies depending on the clonal complex, with CC22 strains having a small type IV SCC*mec* element and CC30 having a large type II SCC*mec* element^42^. Larger SCC*mec* elements contain integrated mobile genetic elements such as plasmids or transposons which can confer resistance to other antibiotics, including erythromycin, kanamycin and tetracycline, and heavy metals, such as mercury and cadmium^42^. However, to the best of our knowledge, this is the first time it has been linked to daptomycin susceptibility.

It has previously been suggested that the presence of a large SCC*mec* element, such as that found in CC30, affects the fitness of the bacteria and compromises their ability to secrete toxins^37,43^. In addition, it has been shown that strains from CC30 are less toxic than those from CC22^31^, possibly explaining our observation that that the CC22 strains were more susceptible to daptomycin than the CC30 strains.

The reason why the presence of a large SCC*mec* element suppresses toxin production is not known, although it has been suggested that PBP2a may cause modifications to the cell wall which interfere with activation of the Agr quorum sensing system^37^. To turn on Agr signalling, the autoinducing peptide must bind to the membrane bound AgrC sensor kinase^44^. It is plausible that alterations in the cell wall could affect this. In support of this, removal of the cell wall with lysostaphin has been demonstrated to increase Agr activity in MRSA strains^37^. We found that expression of *mecA* did not cause any differences in cell wall thickness which could explain the altered Agr activity. However, it is possible that there are other changes within the peptidoglycan, such as altered crosslinking, or differences in the WTA component of the wall.

Daptomycin is often used in combination with a beta-lactam as they have been observed to be synergistic^45–47^. Beta-lactams can increase the toxicity of *S. aureus*^48^, and so it is possible that this increased toxicity could enhance the activity of daptomycin, contributing to their synergistic relationship. By contrast, some antibiotics, including protein synthesis inhibitors such as clindamycin are known to decrease bacterial toxin production^48^. These could therefore promote the inactivation of daptomycin by phospholipids and so may be detrimental to treatment efficacy. The SNAP trial is currently testing whether, amongst other treatment options, the addition of clindamycin to a daptomycin/cefazolin combination improves patient outcomes in MRSA bacteraemia compared to daptomycin and cefazolin^49^.

In summary, we show that PBP2a is a crucial determinant of daptomycin susceptibility. Strains expressing *mecA* produce lower amounts of PSMs, promoting daptomycin inactivation by phospholipids. Therefore, inhibition of PBP2a may be a viable strategy to enhance daptomycin efficacy and improve patient outcomes.

## Materials and methods

The strains used in this study are shown in Table 1. The clinical isolates used are described in Recker *et al*.^*31*^ *S. aureus* was routinely grown on tryptic soy agar (TSA) plates at 37 °C or in tryptic soy broth (TSB) at 37 °C with shaking (180 rpm). Where appropriate, media were supplemented with 10 μg ml^-1^ erythromycin or 5 μg ml^-1^ chloramphenicol. Strains carrying p*itet* or p*itet-mecA* were incubated with 200 ng ml^-1^ anhydrotetracycline.

**Table 1:**
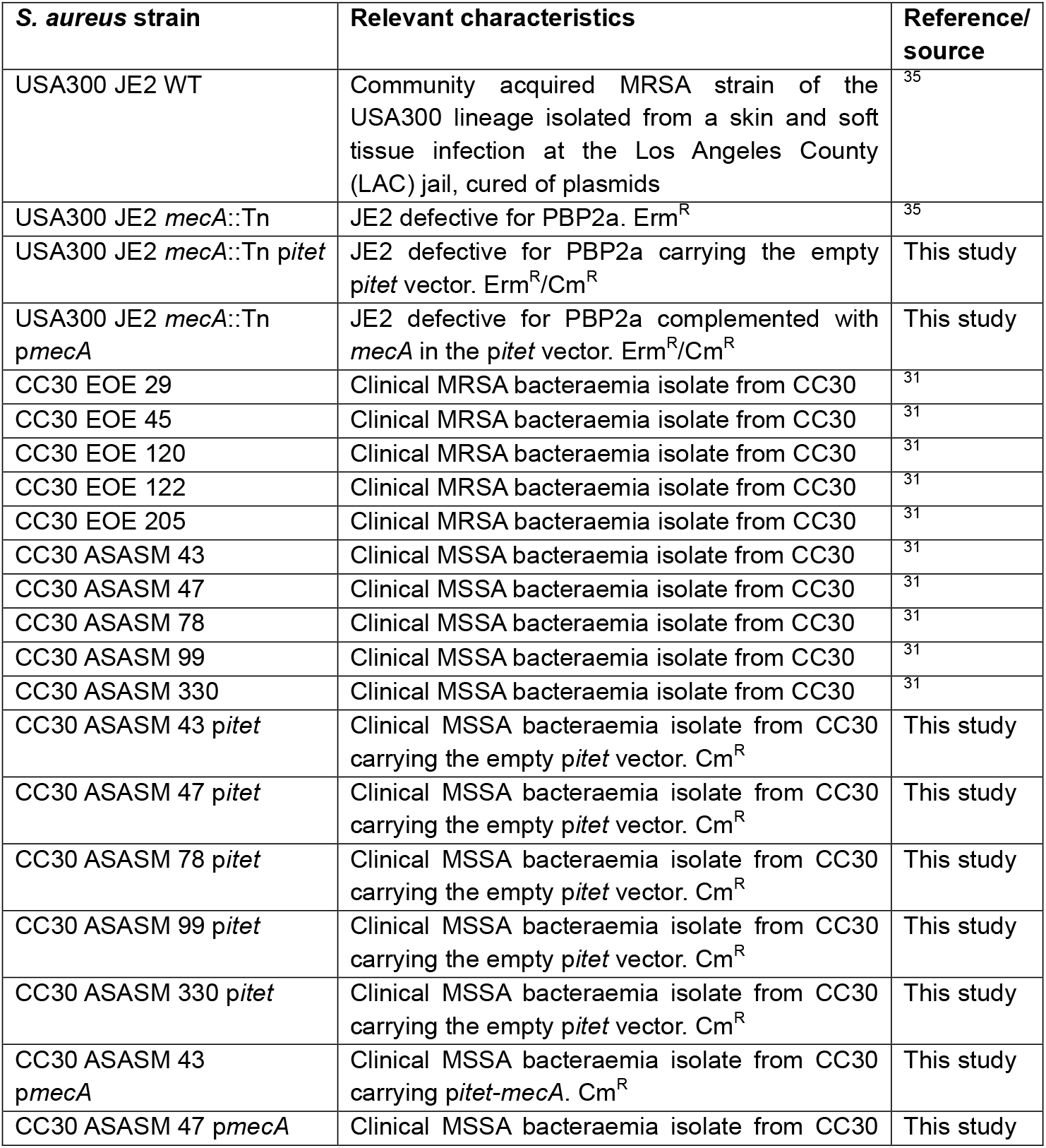
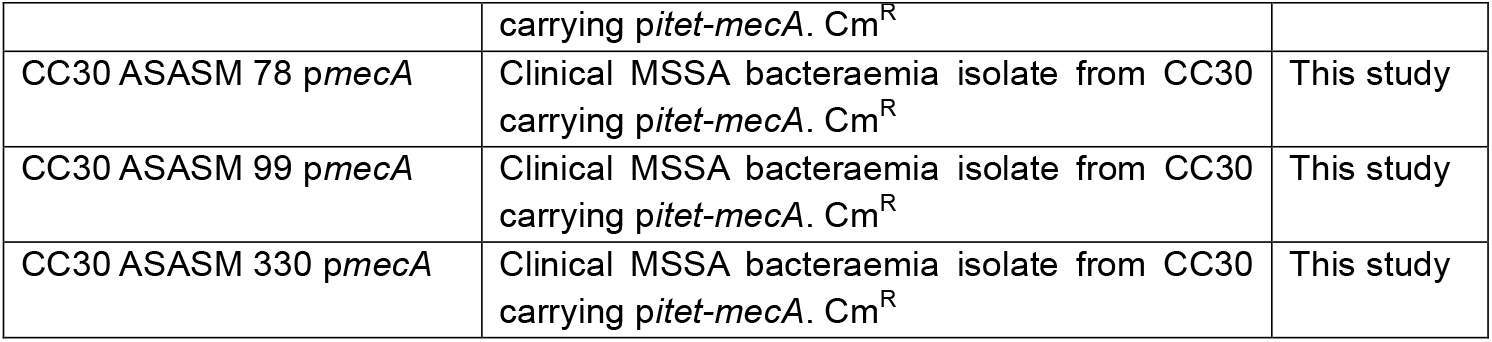
Bacterial strains used in this study.

### Measurements of daptomycin susceptibility

To test the susceptibility of the clinical isolates, strains were grown overnight in 1 ml TSB in bijou tubes at 37 °C with shaking (180 rpm) before being diluted to 10^8^ CFU ml^-1^ in 1 ml TSB supplemented with 1.25 mM CaCl_2_ and 10 μg ml^-1^ daptomycin. Strains were incubated for 2 h at 37 °C with shaking (180 rpm) before survival was determined by 10-fold serial dilutions in PBS and plating onto TSA.

In other cases, strains were grown overnight in 3 ml TSB in universal tubes before being diluted to 10^8^ CFU ml^-1^ in 3 ml TSB supplemented with 1.25 mM CaCl_2_ and 10 μg ml^-1^ daptomycin. Strains were incubated for 6 h at 37 °C with shaking (180 rpm) before survival was determined by 10-fold serial dilutions in PBS and plating onto TSA. Where appropriate, spent culture supernatant was generated by growing *S. aureus* overnight in TSB and removing bacteria by centrifugation and filtration through a 0.2 μm filter. This spent culture supernatant was then supplemented with 1.25 mM CaCl_2_ and 10 μg ml^-1^ daptomycin before bacteria were added at 10^8^ CFU ml^-1^.

### Construction of p*mecA* and complementation of strains

The *mecA* gene was amplified from JE2 WT DNA using the primers avrII_mecA_Fw (5’-AGTCACCTAGGATGAAAAAGATAAAAATTGTTCCAC-3’) and pmeI_mecA_Rev (5’-AGTCAGTTTAAACCACTGTTTTGTTATTCATCTATATCG-3’). It was then digested with avrII and pmeI before being ligated into p*itet* which had been similarly digested using T4 ligase. The plasmid was then transformed into *E. coli* DC10B, before being electroporated into *S. aureus* RN4220 and finally transduced into either the JE2 *mecA*::Tn mutant or the CC30 clinical strains using φ11.

### Determination of surface charge using FITC-PLL

Surface charge was measured as described previously^26^. Briefly, 200 μl aliquots of overnight cultures were washed and resuspended in 200 μl PBS supplemented with 80 μg ml^-1^ FITC-PLL. Samples were incubated at room temperature in the dark for 10 min before being washed three times in PBS, resuspended in 200 μl PBS and the fluorescence determined using a TECAN Infinite 200 PRO plate reader with an excitation of 485 nm and an emission of 525 nm.

### Quantification of cell wall thickness by incorporation of HADA

Overnight cultures of *S. aureus* were generated in TSB supplemented with 25 μM HADA in the dark. Cultures were then washed three times in PBS and the fluorescence determined using a TECAN Infinite 200 PRO plate reader with an excitation of 405 nm and an emission of 450 nm.

### Measurement of membrane fluidity using laurdan

Overnight cultures of *S. aureus* were washed and diluted 10-fold in PBS. Aliquots (200 μl) were incubated in PBS supplemented with 100 μM laurdan for 5 min at room temperature in the dark. Samples were washed four times in PBS before membrane fluidity was determined by measuring fluorescence (excitation 330 nm; emission 460 and 500 nm) using a TECAN Infinite 200 PRO plate reader. The generalised polarisation (GP) value was calculated using the formula: GP = (I_460_-I_500_)/(I_460_+I_500_), where I_460_ and I_500_ are the emission intensities at 460 and 500 nm respectively.

### Extraction and quantification of staphyloxanthin

Aliquots (1 ml) of overnight cultures of *S. aureus* were centrifuged and resuspended in 150 μl methanol. Samples were incubated for 30 min at 42 °C before centrifugation and the absorbance of the supernatant determined at 462 nm.

### Extraction and quantification of WTA

Aliquots (1 ml) of overnight cultures of *S. aureus* were washed with 1 ml 50 mM MES (pH 6.5) (Buffer 1) and resuspended in 10 ml 50 mM MES (pH 6.5) supplemented with 4% SDS (Buffer 2). Samples were boiled for 1 h and centrifuged before being washed twice in Buffer 2, once in 50 mM MES (pH 6.5) supplemented with 2% NaCl (Buffer 3), and once in Buffer 1. The pellet was resuspended in 1 ml 20 mM Tris-HCl pH 8, 0.5% SDS and digested with 20 μg proteinase K for 4 h at 50 °C. The pellet was washed once with Buffer 3 and three times with water, then resuspended in 125 μl 0.1 M NaOH and incubated for 16 h at room temperature. After centrifugation, 100 μl supernatant was neutralised with 25 μl 1 M Tris-HCl (pH 7.8) and analysed by PAGE. 10 μl aliquots of WTA samples were separated on a 20% native polyacrylamide gel by electrophoresis using 0.1 M Tris, 0.1 M Tricine, pH 8.2 running buffer. Gels were then stained with alcian blue (1 mg/ml in 3% acetic acid), destained with water and visualised using a light box. Band intensity was quantified using ImageJ.

### Determination of antibacterial activity of daptomycin

Strains were grown overnight in 3 ml TSB in universal tubes before being diluted to 10^8^ CFU ml^-1^ in 3 ml TSB supplemented with 1.25 mM CaCl_2_ and 10 μg ml^-1^ daptomycin. At each time point, aliquots were taken and centrifuged to remove the bacteria. MHA plates supplemented with 1.25 mM CaCl_2_ were overlaid with 100 μl of the JE2 WT strain at 10^7^ CFU ml^-1^, then wells made in the agar and 100 μl supernatant added. Plates were incubated for 16 h at 37 °C before the diameters of the zones of clearance were measured. TSB supplemented with 1.25 mM CaCl_2_ and 10 μg ml^-1^ daptomycin acted as a positive control and values are presented as a percentage of the positive control or as zone size in mm.

### Determination of haemolysis

Aliquots (100 μl) of supernatants from overnight cultures of *S. aureus* were serially diluted in 2-fold steps in TSB before being mixed with 100 μl 2 % defibrinated sheep blood in PBS. Samples were incubated statically for 1 h at 37 °C before being centrifuged to remove unlysed erythrocytes. Supernatants (100 μl) were moved to a fresh 96 well plate and absorbance determined at 450 nm. Erythrocytes incubated with TSB or TSB supplemented with 1% Triton-X100 were negative and positive controls and values are presented as a percentage of the positive control.

### Measurements of PSM production

Supernatants from overnight cultures of *S. aureus* were analysed by SDS-PAGE using 15% polyacrylamide gels in sample buffer containing bromophenol blue. The gels were stained using Coomassie blue and the PSMs can be visualised as a band which runs ahead of the dye front. Relative band intensity was quantified using ImageJ.

### Statistical analyses

CFU data were log_10_ transformed and presented as the mean ± standard deviation. All experiments consisted of three or more independent replicates and were analysed by t*-*test, one-way ANOVA or two-way ANOVA with appropriate *post-hoc* multiple comparison test using GraphPad Prism (V10.0), as described in the figure legends.

## Acknowledgements

All authors acknowledge the provision of strains by the Network on Antimicrobial Resistance in Staphylococcus aureus (NARSA) Program: under NIAID/ NIH Contract No. HHSN272200700055C.

